# *Admp* regulates tail bending by controlling ventral epidermal cell polarity via phosphorylated myosin localization

**DOI:** 10.1101/2021.09.21.461063

**Authors:** Yuki S. Kogure, Hiromochi Muraoka, Wataru C. Koizumi, Raphaël Gelin-alessi, Benoit Godard, Kotaro Oka, C. P. Heisenberg, Kohji Hotta

## Abstract

The transient but pronounced ventral tail bending is found in many chordate embryos and constitutes an interesting model of how tissue interactions control embryo shape (Lu et al., 2020). Here, we identify one key upstream regulator of ventral tail bending in the ascidian *Ciona* embryo. We show that during early tailbud stage, ventral epidermal cells exhibit a boat-shaped morphology (boat cell) with a narrow apical surface where phosphorylated myosin (pMLC) accumulated. We further show that interfering with the function of the BMP ligand Admp leads to pMLC localizing to the basal instead of the apical side of ventral epidermal cells and a reduced number of boat cells. Finally, we show that cutting ventral epidermal midline cells at their apex using a ultraviolet laser relaxes ventral tail bending. Based on these results, we propose a novel function for Admp in localizing pMLC to the apical side of ventral epidermal cells, which causes the tail to bend ventrally by resisting antero-posterior notochord extension at the ventral side of the tail.

**Summary Statement:** *Admp* is an upstream regulator of tail bending in the chordate *Ciona* tailbud embryo, determining tissue polarity of the ventral midline epidermis by localizing phosphorylated myosin.

## Introduction

Although chordates display diverse shapes and sizes in the adult stage, they have similar shapes during their organogenesis period, called the phylotypic stage (Sander, 1983). During the phylotypic stage, chordates pass through neurulation and subsequently reach the tailbud stage. At the chordate tailbud stage, the embryo tail elongates along the anterior-posterior (AP) axis, and most tailbud embryos become curved with their tail bending ventrally (Richardson et al., 1997).

The ascidian tunicate *Ciona intestinalis* type A *(Ciona robusta)* embryo also shows a curved body shape with its tail bending ventrally (ventroflexion) during the early- to mid-tailbud stages [stage (st.) 19 to st. 22], after which bending relaxes again, and eventually the tail bends dorsally (dorsiflexion). This dynamic body shapes change occur even if the egg envelope is removed, suggesting that *Ciona* tail bending can occur in the absence of external spatial confinement (Hotta et al., 2007; Lu et al., 2020).

During tail extension, notochord cells change their shape by circumferential contraction during st. 21 to 24 (Lu et al., 2019; Mizotani et al., 2018; Sehring et al., 2014). This circumferential contraction, when applied on a compression-resisting system, such as the notochord, is converted into a pushing force along the AP axis of the notochord, thereby elongating the notochord (Lu et al., 2019; Miyamoto and Crowther, 1985).

Although it has been shown that an AP pushing force is exerted by each notochord cell (Sehring et al., 2014; Zhou et al., 2015), and that tail elongation is achieved by the notochord actively producing AP elongating forces (Miyamoto and Crowther, 1985; Sehring et al., 2014; Spemann, 1987; Ubisch, 1939), the mechanism by which the tail bends in the tailbud stage embryo is only incompletely understood. Recently, Lu et al. (2020) have shown that tail bending in *Ciona* during the early tailbud stages (st. 18 to st. 20) is caused by the actomyosin cytoskeleton displaying different contraction forces at the ventral compared to the dorsal side of the notochord. However, the upstream regulators involved in *Ciona* tail bending, and the morphogenetic mechanisms driving tail bending after st. 20 remain unclear.

In this study, we used a combination of genetic, cell biological and biophysical/ three-dimensional (3D) imaging experiments to show that *Admp* regulates cell polarity by determining the localization of phosphorylated myosin (pMLC) at the apex of ventral midline epidermal cells. This ventral epidermal myosin accumulation leads to ventral tail bending by resisting notochord-driven AP tail elongation specifically at the ventral side during mid-tailbud stages.

## Results

### Admp is required for ventral but not dorsal tail bending

Previous studies about Admp function weren’t focused on tail bending but the phenotype is apparent in their images. The knock-down of Admp has been shown to cause reduced ventral tail bending in mid-tailbud stage *Ciona* embryos (Imai, 2006; Imai et al., 2012; Pasini et al., 2006). As these studies did not focus on the tail bending morphant phenotype, we decided to mechanistically dissect how Admp function in ventral tail bending.

To confirm that Admp indeed is required for *Ciona* ventral tail bending, we first performed microinjection of *Admp* morpholinos (MO) and observed the morphant phenotype by recording time-lapse movies (Fig.1A, Suppl. Mov. 1). We found that ventral tail bending (ventroflexion; Fig. 1A, red arrow) was not occurring in *Admp* morphant embryos at the mid-tailbud stages; in contrast, dorsal tail bending (dorsiflexion; Fig. 1A, yellow arrows) was unaffected in morphant embryos. Comparing the bending angle of *Adm*p morphant with wild tye (WT) embryos at st. 18 to 22, when WT ventroflexion occurs (Fig. 1B), showed that the degree of ventroflexion was significantly reduced in morphant embryos (Fig. 1C; N = 11/11). Likewise, embryos treated with dorsomorphin, an Admp/BMP signaling inhibitor, also displayed a significantly reduced ventral tail bending angle (Fig. 1D; N = 5–12, Suppl. Fig. 1A and 1B). Together, these experiments indicate that Admp/BMP signaling regulates the ventroflexion of ascidian tailbud embryos.

**Fig. 1.**
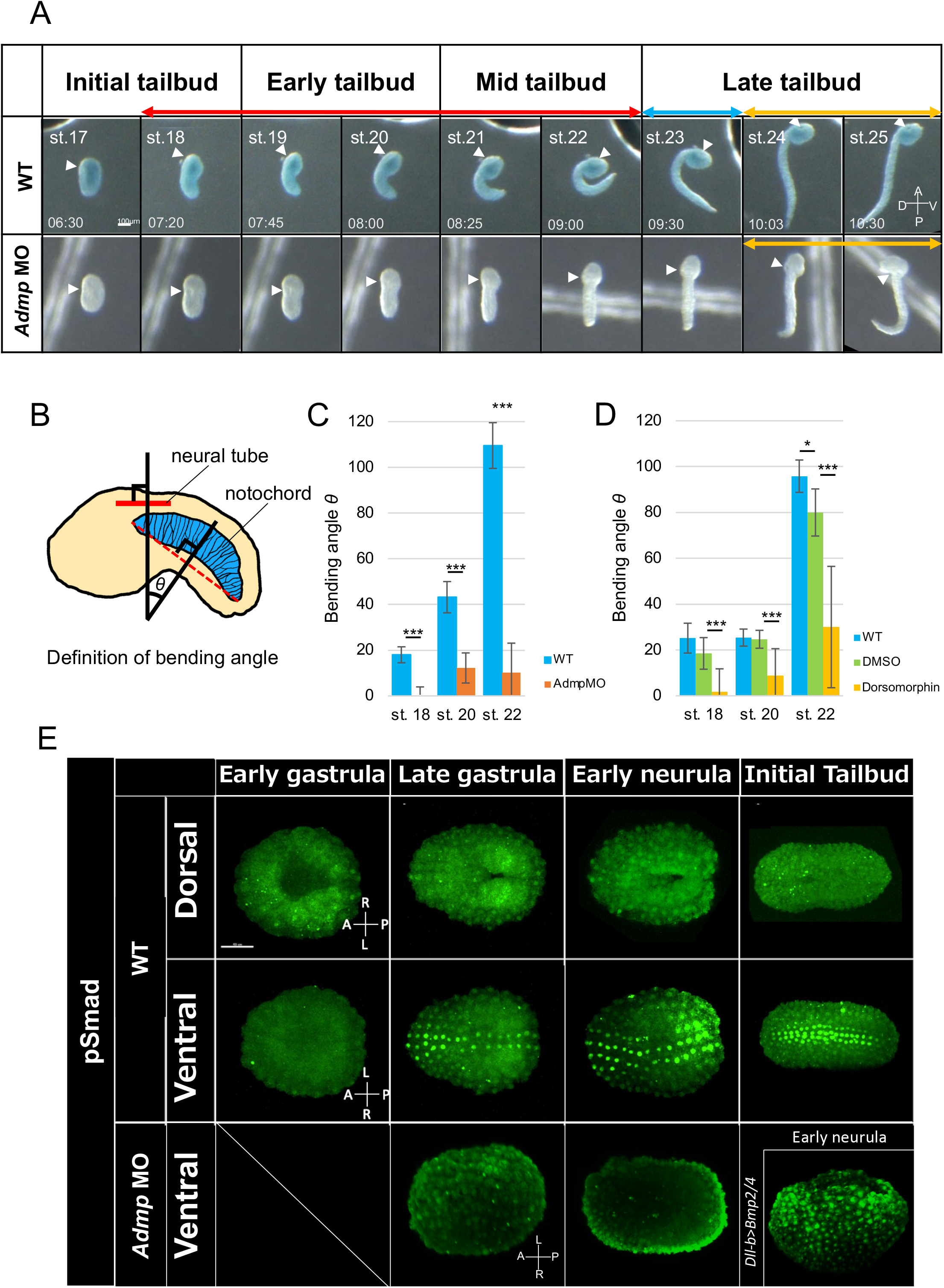
*Admp* affects the tail bending of early tailbud stage embryo, and pSmad was detected at ventral midline epidermis. (A) Time-lapse movie of WT and *Admp* MO embryos (N = 6/6). WT and *Admp* MO embryos were developed in the same dish, and WT embryos were stained by NileBlue B to distinguish them. Note that *Admp* MO suppressed ventral tail bending (“ventroflexion”) during early to mid taibud stages (red double-headed arrow) but not dorsal tail bending (“dorsiflexion”) during the late tailbud stage (yellow double-headed arrow) after “relaxation” (blue double-headed arrow) at the beginning of late tailbud stage. The developmental stage and time after fertilization are shown in each WT picture. Scale bar = 100 μm. A: anterior, P: posterior, D: dorsal, and V: ventral. (B) The definition of the bending angle. The bending angle of the embryo tail was defined as the intersection angle of the straight line perpendicular to the neural tube of the trunk and the anterior-posterior border of the notochord cell, which is the 20th from the anterior side. (C) Quantifying the bending angle θ of WT (n = 8) and *Admp* MO (n = 5~15) embryo at st. 18, st. 20, and st. 22. The bending angle was significantly reduced in *Admp* MO embryos. (D) Quantification of the bending angle θ of WT (n = 8), DMSO-treated embryos (n = 6~7), and Dorsomorphin-treated embryos (n = 9~12). The bending angle was significantly reduced in Dorsomorphin-treated embryos. (E) An antibody staining against phosphorylated Smad1/5/8 (green). The Admp/BMP signals were detected from the late gastrula at ventral midline cells in WT (parenthesis). On the other hand, no signals were detected in *Admp*-MO. Ectopic signals were detected at the whole-epidermal cells in the *Dll-b>Bmp2/4* embryo (right bottom panel). Scale bar = 50 μm.

Admp is a BMP ligand, which in *Ciona*, has been reported to induce the differentiation of ventral peripheral neurons (Imai et al., 2012; Waki et al., 2015), with the homeobox gene *Msxb* functioning as a downstream effector of Admp signaling in this process (Imai et al., 2012). We thus investigated whether *Msxb* might also functions as a downstream effector of Admp signaling in *Ciona* ventroflexion. However, ventroflexion appeared normal in *Msxb* morphant embryos (Suppl. Fig. 1C), suggesting that Msxb, unlike the situation in neuronal differentiation, does not function as a downstream effector of Admp signaling in ventroflexion (Roure and Darras, 2016; Waki et al., 2015).

### Smad phosphorylation in ventral midline epidermal cells

In *Ciona, Admp* is expressed in the endoderm and lateral epidermis (Imai et al., 2012). In vertebrates, *Admp* is expressed first dorsally within the embryo, and then moves to the opposite side to specify the ventral fate, but it is difficult to predict the place of Admp activity from its gene expression pattern. Moreover, *Admp* promotes *bmp4* expression and controls the positioning of *bmp4* expression during regeneration of left-right asymmetric fragments in planarian (Gaviño and Reddien, 2011).

In *Ciona, Admp* expression appears normal in *Bmp2/4* morphants, but *Bmp2/4* expression is suppressed in *Admp* morphants (Imai et al., 2012). Furthermore, the BMP target Smad is phosphorylated by Admp signaling, followed by translocation of phosphorylated Smad into the nucleus and activation of target genes (Blitz and Cho, 2009; De Robertis, 2009; Imai et al., 2012). In line with this, Smad phosphorylation and activation in ventral epidermal cells is reduced in *Ciona Admp* morphant at the late gastrula stage (Fig.1E; Waki et al., 2015).

To determine when and where within the tailbud stage *Ciona* embryo Admp/BMP signaling is activated, we performed antibody staining of phosphorylated pSmad1/5/8 (Fig. 1E). Consistent with a previous studies (Waki et al., 2015), pSmad staining was observed in ventral midline epidermal cells after the late gastrula stage (Fig. 1E), whereas no specific signal was detected in other regions, including notochord, from gastrula to the initial tailbud period. This indicates that Admp/BMP signal is specifically activated in ventral midline epidermal cells.

Asymmetric activation of actomyosin contractility in notochord cells has recently been proposed to be responsible for ventroflexion during st. 18 to st. 20 (Lu et al., 2020). To test whether Admp functions in ventroflexion by affecting asymmetric actomyosin contraction within notochord cells, we analyzed Actin localization in Admp signaling defective embryos. We found that in both dorsomorphin-treated and *Admp* morphant embryos, asymmetric actin localization in notochord cells remained unchanged (Suppl.Fig.2). This indicates that Admp/BMP signaling affects ventroflexion independently from the proposed function of asymmetric actomyosin contraction in notochord cells.

### Admp is required for ordered cell-cell intercalation of ventral epidermal cells

Next, we investigated the dynamics of dorsal and ventral epidermal cell rearrangements during ventral tail bending from st.18 to 24 (Suppl. Fig. 3). Cell-cell intercalation of ventral epidermal cells started at st.19 and was completed by st. 24. The tail epidermis of the ascidian embryo is finally elongated along the AP axis by arranging epidermal cells in a row along this axis through cell-cell intercalation (Suppl. Fig. 3; Hotta et al., 2007). Interestingly, during st. 19 to st. 22, the early phase of epidermal cell-cell intercalation when ventroflexion occurs, intercalation was not associated by an AP elongation of the ventral tail, while during later stages of epidermal cell-cell intercalation from st. 22 to st. 24, intercalation was accompanied by ventral tail elongation (Fig.2AB). We thus hypothesized that epidermal cell dynamics during the early intercalation period contribute to ventroflexion. To test this hypothesis, we compared epidermal cell dynamics between WT and ventroflexion-deficient *Admp* morphant embryos.

**Fig. 2.**
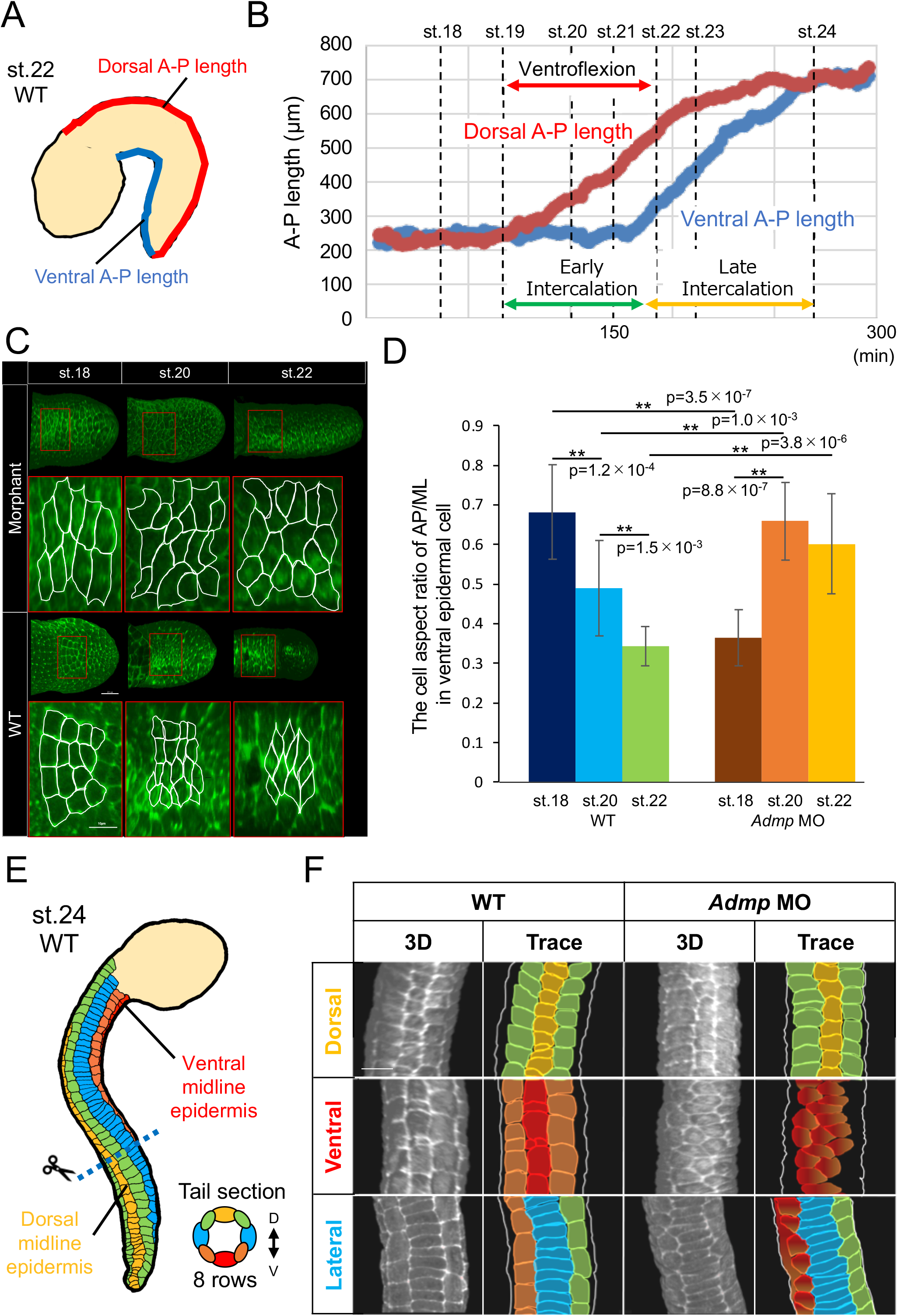
Comparison of the phenotype of *Admp* MO with WT embryo. (A) The measurement of the tail length of the dorsal side (red line) and the ventral side (blue line). (B) Time course of dorsal outline length (red line) and the tail region’s ventral outline length (blue line). Dorsal A-P length of the tail epidermis (red) increases earlier than that of ventral A-P (blue). The ventral A-P length does not change during early intercalation (red double-headed arrow). So, the gap between them is increasing in this period. The ventral A-P length increased at late intercalation (double-headed arrow), and the gap decreased. Each dotted line indicates the corresponding developmental *Ciona* stage. At stage 24, the dorsal A-P and ventral A-P have the same length. (C) The cell shape change of anterior ventral epidermis of WT and *Admp* morphant during early intercalation. The figures show the ventral view. The F-actin was stained by phalloidin. The cell shape was traced with a white line. In all figures, the anterior ventral tail epidermal cells were perpendicular to the direction of observation and enclosed a scale bar: of 20 μm. (D) The AP/ML aspect ratio of the ventral epidermis of WT and *Admp* morphant during early intercalation in Figure 2C was measured with reference to the rectangle to AP-ML direction (WT; st.18 n = 14, st.20 n = 19, st.22 n = 10; *Admp* MO; st.18 n = 10, st.20 n = 16, st.22 n = 16). Asterisk indicates statistically significant (t-test, *: p< 0.05). The error bar indicates SD. (E) The schematic alignment of the tail epidermal cells of WT embryo at st. 24. The cell-cell intercalation of the tail epidermis has finished at st. 24. The tail epidermal cells consist of eight rows: dorsal (yellow), two dorsal medio-lateral (green), ventral (red), two ventral medio-lateral (orange), and two lateral (blue) rows. (F) The alignment of the tail epidermal cells of WT and *Admp* MO embryo at st. 24. There is a specific inhibition of the intercalation of the ventral rows (red and orange) in the *Admp* MO embryo (N = 4 in WT and 4 in *Admp* MO). These mixed colors indicate that we cannot distinguish ventral and ventrolateral epidermal cells, scale bar: 10 μm.

During st.20 to 22 the ventral epidermis in WT embryos showed a preferential accumulation of junctional F-actin in the medio-lateral direction (ML accumulation) (Fig. 2C). Antibody staining of pMLC also showed such ML accumulation, especially at st. 19 to 22 (Suppl. Fig. 4). In contrast, no such ML accumulation was observed in *Admp* morphant embryos during st. 20 to 22 (Fig. 2C). In addition, while the AP/ML aspect ratio of ventral epidermal cells decreased in WT embryos during st. 18 to 22, no such decrease was found in *Admp* morphants (Fig. 2D). This suggests that *Admp* is required for proper asymmetric junctional actin accumulation and ML elongation of ventral epidermal cells during early intercalation. At st. 24, the tail epidermis in WT embryos became organized into eight distinct single-cell rows as a result of cell-cell intercalations (Fig. 2EF) (Hotta et al., 2007; Pasini et al., 2006). Moreover, these eight rows, consisting of three rows of dorsal, two rows of lateral, and three rows of ventral epidermal cells, were closely aligned (Fig. 2F, WT). In contrast, the ventral three-rows in *Admp* morphant embryos were disorganized into one or two rows, making it difficult to clearly distinguish between midline and medio-lateral cells (Fig. 2F, *Admp* MO; mixed orange/red color). Dorsomorphin-treated embryos showed a similar disordered ventral midline intercalation phenotype (Suppl. Fig. 5), further supporting the notion that *Admp* regulates ordered ventral epidermal cell-cell intercalation.

### Ventral epidermal cells display a ‘boat-like’ morphology during ventroflexion

We suspected that defective ventroflexion in *Admp* morphant embryos involves changes in ventral epidermal cell morphologies (Fig. 2F). To further investigate what detailed morphological change occurs in the ventral epidermal cells during this period, we monitored changes in single ventral epidermal cell morphology by 3D imaging. This revealed that ventral epidermal cells acquire a distinctive ‘boat-like’ morphology (boat-cell), characterized by a larger area on the basal surface (Fig. 3AB, yellow areas) as compared to the apical surface (Fig. 3AB, red areas), and ridges at both ends oriented along the ML direction. Almost all anterior ventral epidermal cells showed this shape (Suppl. Mov. 2), consistent with previous reports that tail bending only occurs in the anterior tail of *Ciona* (Lu et al., 2020). The shape of boat-cell is characterized by a triangular-shaped cross-section where the apical surface is entirely constricted (triangular-shaped section of boat-cell; TSBC), and a square-shaped cross-section (square-shaped section of boat-cell; SSBC), where some apical surface is left (Fig. 3A, B). In *Admp* morphant embryos at st. 22, the number of TSBCs in ventral epidermal cells was strongly reduced (Fig. 3D and 3E; *Admp* MO, N= 12, WT, N = 7, p = 0.05×10^-5), while the number of non-TSBCs was increased (the section of non-boat cell) indicative of a reduced number of boat-cells in all ventral epidermis sections of morphant embryos (Suppl. Mov. 2).

**Fig. 3.**
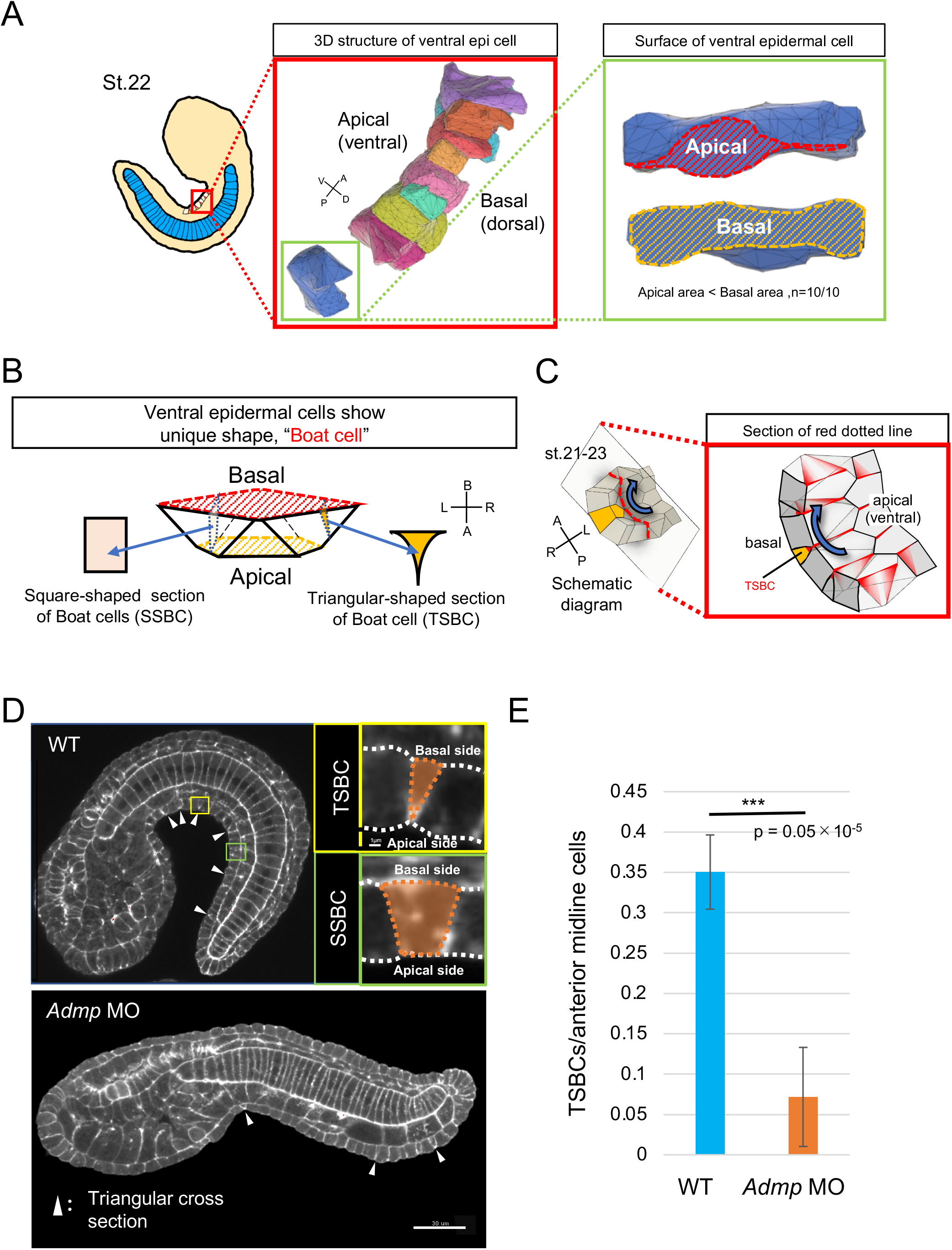
Three-dimensional reconstruction of ventral midline epidermal cells during tail bending. (A) The 3D model reflecting the shape of each midline ventral epidermal cell at st. 22 was reconstructed using Avizo6 software (inside of red rectangular). Representative morphology of ventral midline cells is shown (inside the green rectangle). Each cell is bipolar laterally and has protrusions from basal parts. Note that the apical area (red) is smaller than the basal area (yellow). (B) The 3D model reflects the shape of ventral epidermal cells in (A). The ventral epidermal cells show a distinctive shape, a “boat cell”. The sections of the boat cell show triangular-shaped (TSBC) or square-shaped (SSBC) depending on the plane. (C) Schematic drawing of the cell-cell intercalation of the ventral midline epidermal cells using boat cells during the tail bending period. The ventral midline epidermis consists of boat cells with a smaller apical domain area than the basal domain; the overall tissue becomes bending (blue arrow). The orange cell’s midline section (cut in the red dotted plane) shows TSBC. The orange cell starts intrusion from basolateral sides with pMLC protrusions (red-colored). The 3D models were generated by FUSION 360 educational ver. (Autodesk). (D) Midline section view of WT tailbud embryo and *Admp* MO embryo at st. 22 by staining of F-actin. Arrowheads indicate the position of TSBC. The regions in a yellow rectangle are enlarged, showing a TSBC. The regions in a green rectangle are enlarged, showing an SSBC. (E) The ratio of the number of TSBC and the number of midline cells. It has been reported that the driving forces of the tail bending originate in the anterior part of the tail (Lu et al., 2020). Therefore, we counted the number of TSBC in the anterior part of the tail. TSBCs were significantly reduced in *Admp* MO embryos. Asterisks indicate statistical significance (t-test, *: p< 0.05). The error bar indicates SD (WT, N = 8; Admp MO, N = 6).

### Admp/BMP signaling is required for the localization of the pMLC to the apical side of ventral epidermal cells

To understand how this distinctive boat-cell morphology (Fig. 3A and 3B) arises, we performed both F-actin/Phalloidin staining and antibody staining for pMLC. This showed an accumulation of both F-actin and pMLC at the apical side of TSBC (Fig. 4A, WT arrowheads, 4B). Interestingly, the localization of pMLC to the apical side was significantly decreased in *Admp* morphant embryos (Fig. 4C; *Admp* MO, n = 7, WT, n = 9, p = 0.01), suggesting that Admp triggers the formation of TCBCs, and thus boat-cell shape, by localizing pMLC to the apical side of ventral midline cells.

**Fig. 4.**
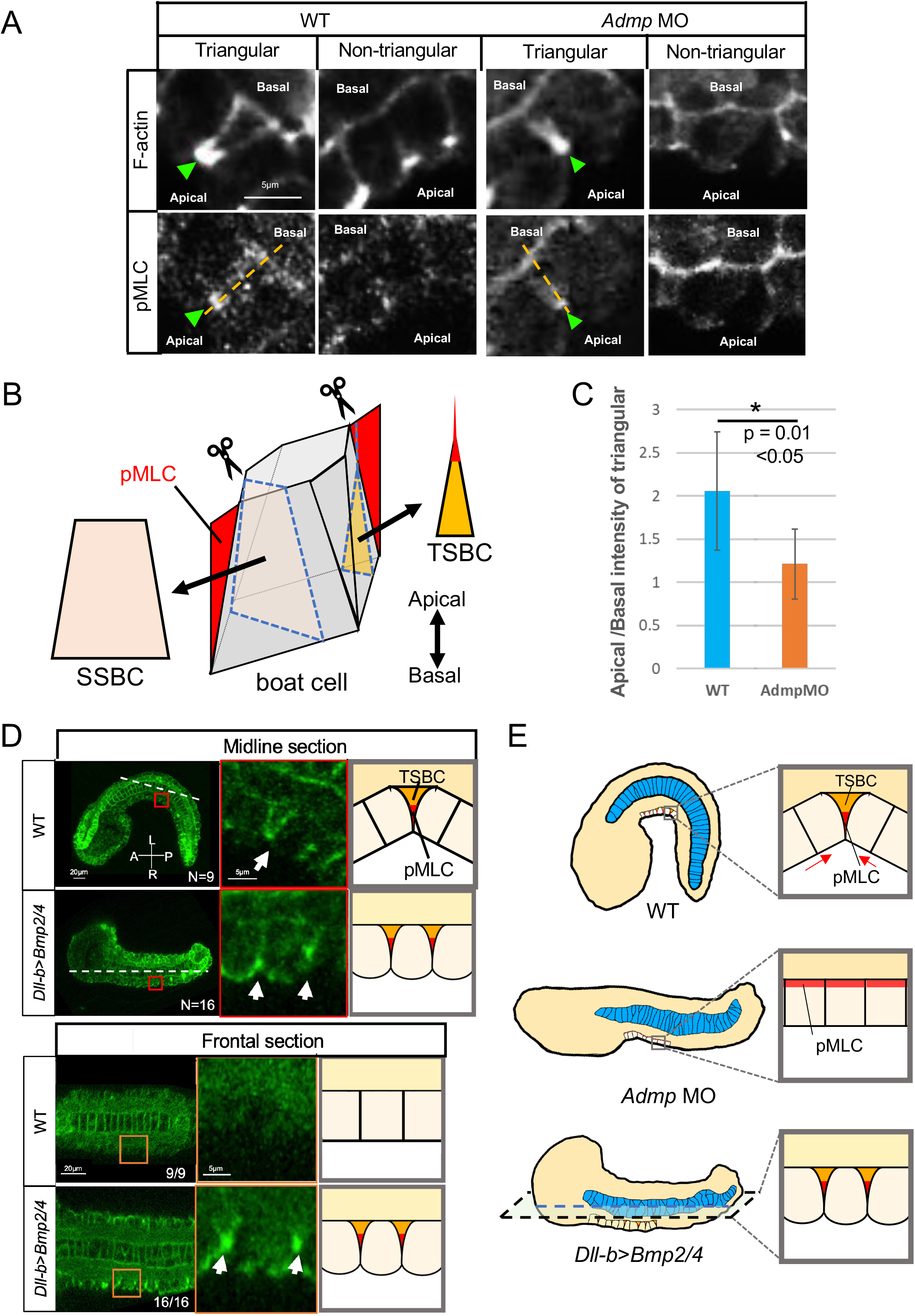
The distribution of pMLC of tail epidermis depends on the Admp/BMP pathway. (A) Double staining of F-actin and phosphorylated myosin light chain (pMLC) of the ventral midline of WT and *Admp* MO embryo at st. 22. Both F-actin and pMLC accumulated at the apical side of triangular cells (arrowheads). The loci of the measurement of the relative pMLC intensity ratio between the apical and basal side of the epidermis in WT and *Admp* MO(B) are shown as yellow lines. (B) The 3D model of a boat cell reflecting the localization of pMLC (red colored). Different section planes show different shapes, triangular-shape, TSBC, and square-shape, SSBC. (C) Relative pMLC intensity ratio between the apical and basal side of the epidermis in WT (N = 10, n = 10) and *Admp* MO (N = 7, n = 7). Asterisks indicate statistical significance (t-test, *: p<0.05, **: p<0.01). The error bar indicates SD. (D) The distribution of pMLC in WT and *Dll-b>Bmp2/4* at st. 22. In the midline section, the pMLC was distributed at the apical TSBCs of the ventral midline epidermis in both WT and *Dll-b*>*Bmp2/4* (arrows in red rectangles). On the other hand, in the frontal section, the apical TSBCs of the lateral epidermis were observed in *Dll-b>Bmp2/4* embryo (arrows inside the orange rectangle), but there was no signal of pMLC in the lateral epidermis in WT. The white dotted lines indicate the section plane of the frontal sections. (E) The schematic of the difference in the distribution of pMLC (red) among WT, *Admp* MO, and *Dll-b>Bmp2/4* embryos. The orange-colored cell sections are TSBC.

To test whether Admp/BMP signaling can ectopically affect the localization of pMLC and thereby generate TSBC (Fig. 4D), we performed ectopic *BMP*-expression experiments. In WT embryo, both apical pMLC accumulation and TSBCs were not observed in epidermal cells except ventral epidermal cells, where also pSmad signal was detected (Fig. 4D, a frontal section of WT). In contrast, in embryos ectopically expressing BMP, pSmad signal was detected in all epidermal cells, accompanied by apical pMLC accumulation and TSBC formation not only in ventral tail epidermal cells but also within the remainder of the tail epidermis (Fig. 4D).

These suggests that Admp/BMP signaling is sufficient to induce the localization of pMLC to the apical side of epidermal cells (Fig. 4E), leading to the formation of boat-cells. This cell shape change, again, might resist tail elongation at the ventral tail region, leading to ventroflexion.

### Cutting ventral epidermal cells relaxes ventroflexion

To investigate whether the ventral epidermis indeed locally resists tail elongation, eventually leading to ventroflexion, we cut either ventral or dorsal epidermal cells at their apex along the AP axis using an ultraviolet (UV)-laser cutter (Fig. 5, yellow lines). When dorsal midline epidermal cells were cut, no effect was observed. However, cutting ventral midline epidermal cells led to a strong relaxation of ventroflexion, indicative of stress-release along with the AP axis at the ventral midline epidermal cells (Fig. 5A and 5B, Suppl. Mov. 3). These findings suggest that apically accumulated pMLC in boat-cells generate AP stress in the ventral midline epidermis, resisting tail elongation and thus enabling ventroflexion.

**Fig. 5.**
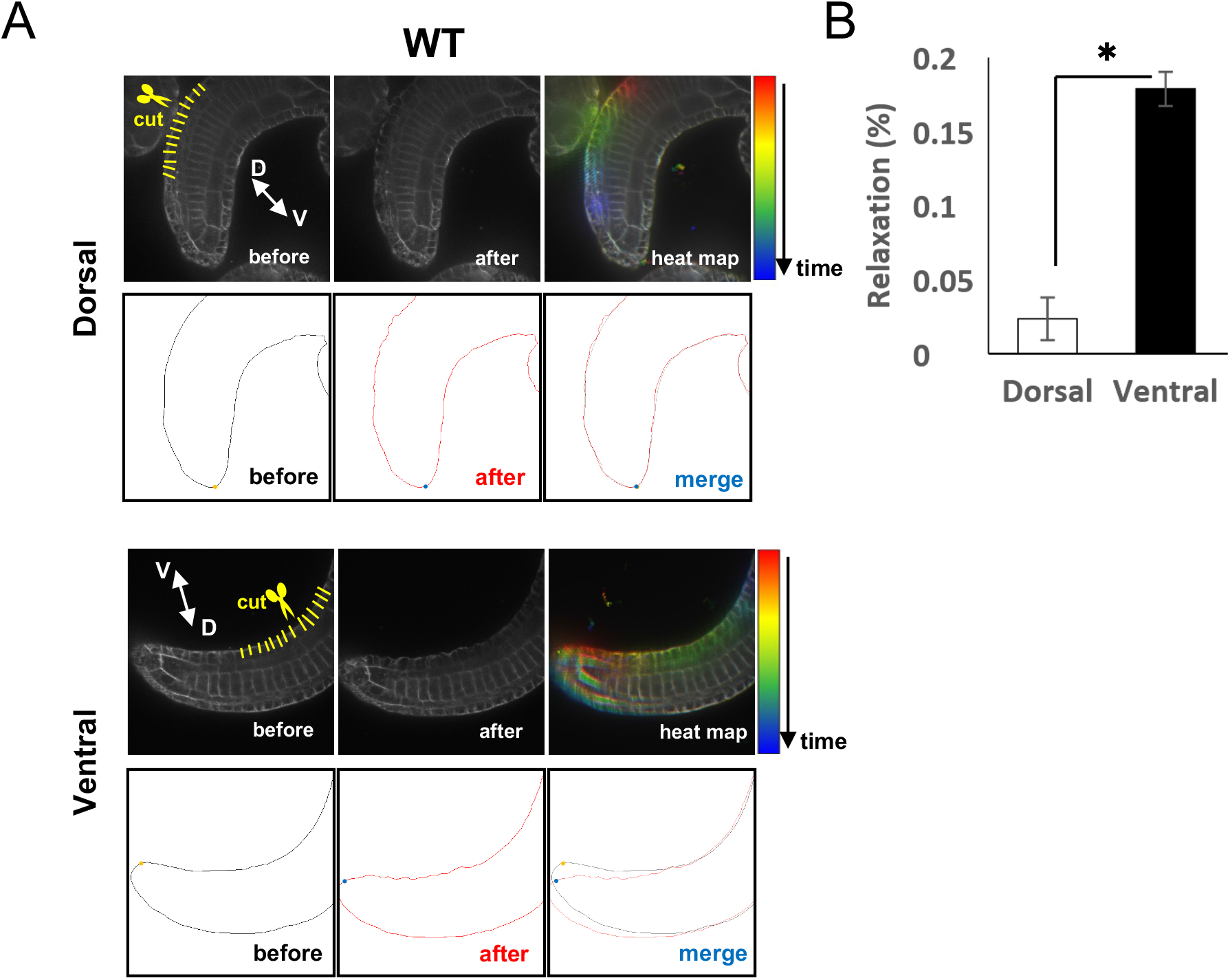
Laser-cut experiment for the AP cell-cell border of the tail epidermis. (A) Laser-cutting of the wild type’s dorsal and ventral midline epidermis at st. 22. Epidermal cells are a monolayer. We cut each cell in an apico-basal direction (yellow lines). Cell membranes were stained by FM4-64. The color bar indicates the time after laser cut from 0 (before) to 30 (after) s post cut. (B) The movement of the area before and after the laser cut was calculated as relaxation (N = 5). Asterisks indicate statistical significance (t-test, *: p< 0.05). The error bar indicates SD.

We further investigated whether the relaxation of ventroflexion by UV-laser cutting depends on whether the cuts were oriented along the AP axis or ML axis in ventral midline cells. This showed that the relaxation of ventroflexion was slower in AP cuts when compared to ML cuts (Suppl. Mov. 4), consistent with ML cuts more efficiently interfering with the AP stress in the ventral epidermis midline than AP cuts.

### pMLC localization predicts development of boat-cell morphology

Next, we asked how boat-cells change their shape during the ventral tailbud period (Fig. 6A). To this end, we analyzed the subcellular distribution of pMLC as proxy of actomyosin contraction in ventral midline cells (ventral view in Suppl. Fig. 4). This revealed that pMLC first emerged at the ML junction of ventral midline epidermal cells at st. 21 (Fig. 6A, st. 21 arrowheads), and localized to the apical side of TSBCs at st. 22 (Fig. 6A, st. 22 arrowheads). At st. 23, the trapezoid shape became apparent at the midline plane (Fig. 6, st. 23; Suppl. Fig. 4), and finally, the apical accumulation of pMLC disappeared at st. 24 (Fig. 6, st. 24; Suppl. Fig. 4). These observations suggesting that TSBC formation is driven by ML contraction as follow. The ML directed localization of pMLC and the cell shape change to boat-cell correspond with ventroflexion, and the pMLC disappearance and the cell shape change to disengage the boat-like morphology correspond with relaxation.

**Fig. 6.**
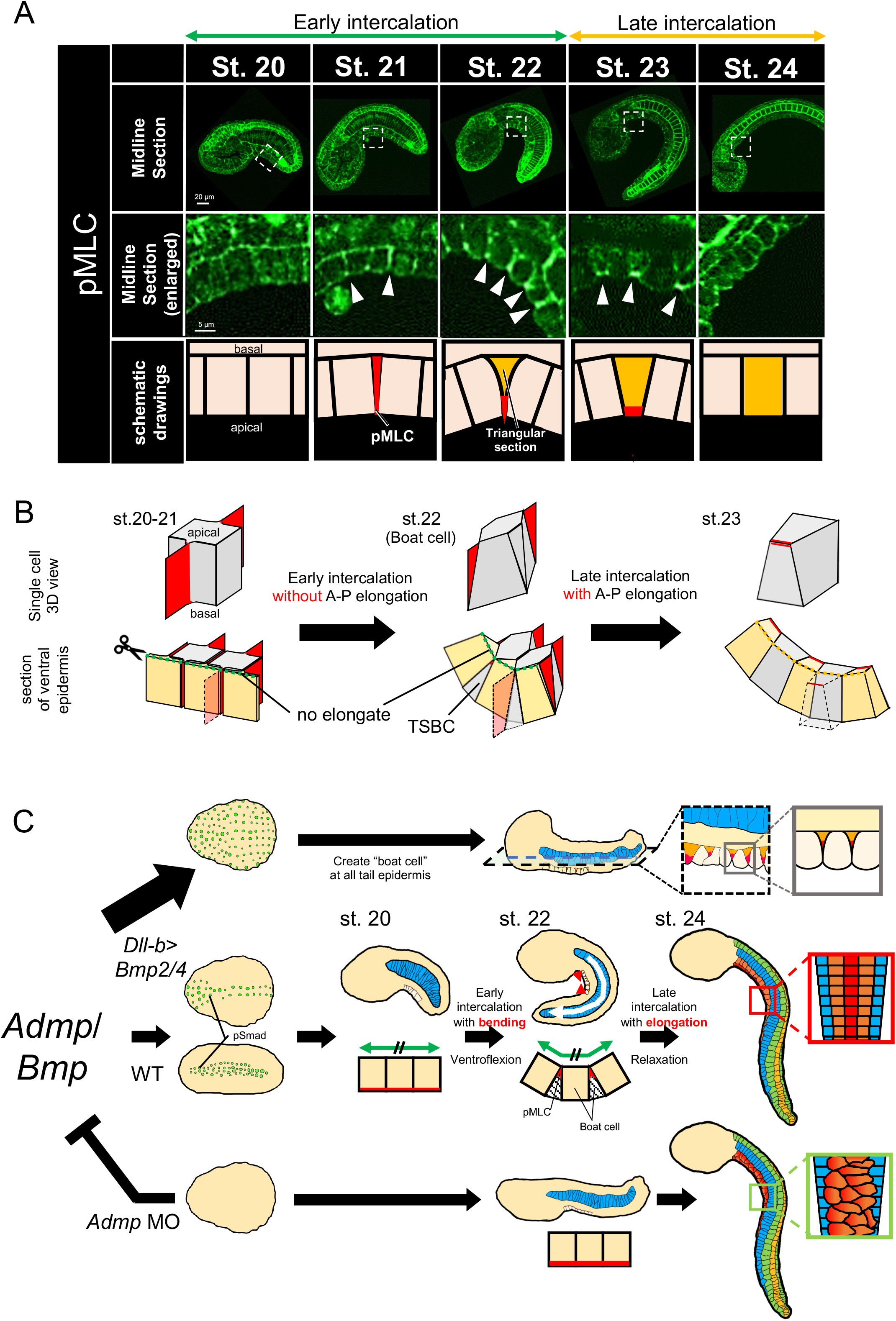
Relationship between the change of the distribution of pMLC and the tail bending during intercalation of midline epidermal cells. (A) (top) The antibody staining of 1-phosphorylated-myosin (pMLC) from st. 20 to 24 in WT. Scale bar = 50 μm. (middle) The enlarged view of the dotted rectangle in each stage. Arrowheads show pMLC accumulation in the apical domain of the ventral midline epidermal cells. (bottom) The schematic drawing of the distribution of pMLC (shown in red). In st. 20, pMLC localized the basal side of the epidermis. In st. 21, pMLC has appeared at the AP cell border of ventral midline epidermal cells. At st. 22, the pMLC accumulated at the apex of the apical domain and the cell shape changed into TSBC of the boat cell in Figure 3C (orange-colored cell). The pMLC localization at the basal side was reduced. At st. 23, the apex of the apical domain becomes broader. At st. 24, the pMLC asymmetrical distribution disappears. (B) The schematic model explains the halt of the AP elongation during the early intercalation period. At the beginning of the intercalation, the total AP length of the midline cells was shown as a green arrow (st. 20). During early intercalation, the ventral epidermal cells change cell shape to become boat cells. The boat cells start to intrude on each other from the basal plane. Still, the total AP length of the apical side was not changed (green arrow) because apically accumulated pMLC (st. 22) resists the AP elongation force of the notochord. In late intercalation, the boat cell intercalates the basal side and the apical side, and the apical area increases, allowing the elongation of AP length (yellow arrow) of the apical side (st. 23). (C) Model of embryonic tail bending in *Ciona.* Admp/BMP signaling (green dots) transmits the signal to the ventral midline epidermal cells as phosphorylated Smad from neurula to the initial tailbud. pSmad translocates the localization of the pMLC from the basal side (dorsal side) to the apical side (ventral side), which changes the cell polarity and promotes the cell-cell intercalation of the ventral midline epidermal cells during early to mid-tailbud stages (st. 20 to st. 23). The ongoing mediolateral intercalation at the ventral epidermis confers a resistance (red arrowheads) to AP elongation force (white arrow) that is possibly provided by the notochord, which causes the bending tail shape in the *Ciona* tailbud embryo at st. 22. Conversely, the *Admp* MO or dorsomorphin treatment disrupts the cell polarity and causes the no tail-bending embryo at st. 20–23 and incomplete intercalation at st. 24 (the flow by the green rectangle). The final overall embryo shape at st. 24 is similar among WT and *Admp* MO embryos, but the ventral cell-cell intercalation was disrupted (red and orange gradation-colored cells). Thus, Admp/BMP signaling, apart from its known role in peripheral nervous system (PNS) differentiation, regulates temporal tail bending during early to middle tailbud stages (st. 20 to st. 23).

## Discussion

### Admp is an upstream regulator of tail bending

Originally, the Admp/BMP pathway was identified as central in establishing, maintaining, and regenerating the DV axis among bilaterian animals (Gaviño and Reddien, 2011). In the ascidian, both gain- and loss-of-function experiments demonstrated that *Admp* expressed in the B-line medial vegetal cells acts as an endogenous inducer of the ventral epidermis midline (Pasini et al., 2006). *Admp* is required for sensory neuron differentiation of the ventral epidermis via the *Tbx2/3* and *Msxb* genes (Waki et al., 2015). Our finding that tail bending was not regulated by *Msxb* (Suppl. Fig. 1C) suggests that Admp functions in ascidian tail bending through different effector pathway(s) than those implicated in ventral epidermal cell fate specification. Admp controls ventral, but not dorsal tail bending by determining early ventral epidermal cell intercalation (Fig. 2), and the shape of ventral epidermal midline cells (Fig. 3) through the localization of pMLC (Fig. 4). Importantly, this does not exclude that genes other than Admp act on tissues other than the ventral epidermis to control ventroflexion.

### Admp controls ordered cell-cell intercalation of ventral epidermal cells

We divided the intercalation period into two phases: during early intercalation, the ventral epidermis does not elongate in an AP direction; during late intercalation, in contrast, the ventral epidermis elongates. Interestingly, ventral tail bending (ventroflexion) occurs during early intercalation, with ventral epidermal cells displaying a flat cell shape elongated along the ML axis (Fig. 2C, D). We assume that this ML cell elongation is caused by the accumulation of actomyosin in ML-oriented protrusion-like extensions of ventral epidermal cells (Fig. 6B) extending cells along the ML axis (Suppl. Fig. 7). No such actomyosin localization and ML elongation was found in *Admp* morphant embryos, suggesting that *Admp* is required for ventral epidermal cell polarization and protrusion formation.

When ventral epidermal cell intercalation is completed (st. 24~), ventral epidermal cells drastically change their polarity into the AP direction in WT embryos. This does not occur to the same extent in *Admp* morphants (Fig. 2D), suggesting that Admp is also required for this later shift in cell polarity.

Ventral tail epidermal cells in WT, but not *Admp* morphant embryos, arrange into three ordered rows at st. 24. This suggests that *Admp* may be required for the cell autonomous ML intercalation of ventral epidermal cells by controlling ML cell polarization and protrusion formation. Notably, in *Admp* morphants, the ventral epidermis was disordered but kept a three-cell width, suggesting that some intercalation of ventral epidermal cells might still occur in the absence of Admp.

How does the intercalation of ventral midline cells contribute to ventroflexion? The ventral epidermis undergoing early cell intercalation does not elongate along the AP axis from st. 20 to st. 22, different from the already fully intercalated dorsal epidermis (Fig. 2B). Assuming that the notochord functions as the main force-generating structure driving tail elongation (Dong et al., 2011; Hara et al., 2013; Lu et al., 2019), the lack of ventral epidermis elongation during st. 20 to st. 22 might locally resist the global notochord-mediated tail elongation, thereby causing the tail to bend ventrally. The lack of ventral epidermis elongation along the AP direction during early intercalation is likely due to the Admp-dependent polarization of ventral epidermal cells along the ML direction. This cell polarization perpendicular to AP direction will limit epidermal AP elongation during intercalation, which again, by resisting global notochord-driven tail elongation, leads to ventroflexion.

### Admp regulates ventral epidermal cell shape changes

Ventral epidermal cells take a distinct boat-cell shape, which likely contributes the ventroflexion (Fig. 4). The preferential accumulation of pMLC in ventral epidermal cells along ML junctions (Suppl. Fig. 4Db’, Eb’) is found at the apical side of cell boundaries of TSBC and/or SSBC and might correspond to protrusion-like extensions formed between interdigitating boat cells (Fig. 6B). The lack of such polarized distribution of pMLC suggests that Admp might be required for both planar and apicobasal polarization of these cells.

As the cell-cell intercalation of the boat cell progresses, the shape of these cells changes from TSBC to a trapezoid shape (Fig. 6A). This shape change occurs during the late intercalation period (Fig. 2B) and leads to ventral epidermis elongation along the AP axis (Fig. 6B, green dotted line). Thus, ventral epidermal cell elongation along the ML axis during early intercalation locally resist notochord-mediated tail elongation, thereby triggering ventroflexion (Fig. 6B, st. 20 to 22). During later intercalation, in contrast, the ventral epidermal midline cells enlarge their apical area and elongate along the AP axis, thereby relaxing the local resistance against tail elongation (Fig. 6B, yellow dotted line st. 23).

How does Admp/BMP signaling regulate both the cell-cell intercalation and the apico-basal polarity of the ventral epidermal cells? Our finding suggest that Admp is required for the preferential localization of pMLC not only at ML junctions between intercalating cells (Suppl. Fig. 4) but also at the apical side by pSmad signaling (Fig. 4D). Recent studies show that SMAD3 drives cell intercalation underlies secondary neural tube formation in the mouse embryo (Gonzalez-Gobartt et al., 2021). Moreover, the BMP-Rho-ROCK1 pathway is thought to target MLC to control actin remodeling in fibroblasts (Konstantinidis et al., 2011). Finally, BMP regulates cell adhesion during vertebrate neural tube closure and gastrulation (von der Hardt et al., 2007; Smith et al., 2021). Yet, how BMP/Smad signaling regulates the localization of the pMLC in ventral epidermal cells is still unclear.

### Model of ascidian ventroflexion

Our findings demonstrate that Admp is required for ventroflexion of the ascidian tail during tailbud stages (st. 18 to st. 22). We propose that Admp phosphorylates Smad in the ventral epidermis. pSmad, in turn, allows early cell intercalation within the ventral epidermis by controlling the localization of the pMLC leading to ventral epidermal cells taking a boat-cell-like shape. This cell shape change limits ventral epidermis elongation along the AP axis, thereby locally resisting global notochord-driven tail elongation causing the tail to bend down (Fig. 6).

The notochord has recently been proposed to display asymmetric contraction forces before st. 20 by the asymmetrical localization of actomyosin in notochord cells (Lu et al., 2020). However, *Admp* morphant embryos displaying straight tails still have ventral bias in notochord actomyosin localization (Suppl. Fig. 2). This suggests that *Admp* is not required for asymmetrical notochord actomyosin localization, and that this asymmetric localization is not sufficient to cause ventroflexion. One possibility is that the ventral accumulation of actomyosin in the notochord might be involved in earlier morphogenetic events, such as notochordal cell intercalation, giving rise to a transient ventral groove (Munro and Odell, 2002).

### The evolutionary roles of *Admp*

Our study provides insight into the molecular and mechanical mechanisms underlying conserved shape changes of chordate embryos, such as tail bending.Tail bending in the tailbud stage embryos is a still understudied morphogenetic process although many genes, including *Admp*, with a critical function in tail bending have been identified [zebrafish: (Esterberg et al., 2008; Willot et al., 2002); frog: (Dosch and Niehrs, 2000; Kumano et al., 2006)]. In invertebrate non-chordate animals, such as sea urchins and hemichordates, Admp is expressed within the embryonic ectoderm (Chang et al., 2016; Lowe et al., 2006). It would thus be interesting to investigate whether the regulation of pMLC subcellular localization by Admp is conserved in primitive chordate embryogenesis and causes the body shape change in these animals.

## Materials and methods

### Ascidian samples

*C. robusta (C. intestinalis* type A) adults were obtained from Maizuru Fisheries Research Station (Kyoto University, Kyoto, Japan), Onagawa Field Center (Tohoku University, Sendai, Japan), and Misaki Marine Biological Station (University of Tokyo, Tokyo, Japan) through the National Bio-Resource Project (NBRP, Japan) and Roscof Marine Station (Roscof, France). Eggs were collected by dissection of the gonoducts. After artificial insemination, fertilized eggs were incubated at 20°C until fixation or observation. Developmental stages followed Hotta’s stages (Hotta et al., 2007; Hotta et al., 2020). In inhibiting phosphorylation of 1P myosin, Y27632 (nacalai tesque, Japan) was applied to embryos at 10 μM after 7 hpf (late neurula, st. 16). Dorsomorphin (Sigma-Aldrich) was applied to embryos at 10 μM after fertilization.

### Immunostaining and quantifying pMLC intensity

To detect activation of the Admp/BMP signaling pathway, we followed the same method described previously (Waki et al., 2015). The signal was visualized with a TSA kit (Invitrogen) using horseradish peroxidase-conjugated goat anti-rabbit immunoglobulin-G and Alexa Fluor 488 tyramide.

The method of pMLC antibody staining was as follows. Embryos were fixed in 3.7% formaldehyde in seawater for 30 min and then rinsed with phosphate-buffered saline with Tween (PBST; 0.2% Triton-X in PBS) for 3 h. Embryos were incubated in PBST containing 10% goat serum for 3 h at room temperature or overnight at 4°C. The primary antibody (anti-rabbit Ser19 phosphorylated-1P-myosin, Cell Signaling, USA) was diluted at 1:50 and then incubated for 3 h at room temperature or overnight at 4°C. The primary antibody was washed with PBST for 3 h. A poly-horseradish peroxidase (HRP) secondary antibody (goat anti-rabbit IgG, Alexa Fluor 488 Tyramide SuperBoost Kit, USA) was applied for 3 h and washed in PBST for 3 h. Alexa Fluor dye tyramide (Alexa Fluor 488 Tyramide SuperBoost Kit) was added to the reaction buffer for 5 to 8 min to induce a chemical HRP reaction. Embryos were dehydrated through an isopropanol series and finally cleared using a 2:1 mixture of benzyl benzoate and benzyl alcohol.

pMLC accumulation was quantified by measuring the intensity along the ventral tail epidermis using Fiji image analysis software. The signal in the brain region was taken as the positive control because its signal was detected even in Y27632 treated embryos, indicating RhoA kinase (ROCK)-independent expression. The relative intensity of pMLC normalized to the intensity of the brain region in each individual was calculated by ImageJ for the comparative analysis among different individuals.

### Laser-cutter experiment

UV-laser cutting experiments were performed on tailbud *Ciona* embryos. An inverted Axio Observer Z1 (Zeiss) microscope equipped with a confocal spinning disk (Andor Revolution Imaging System, Yokogawa CSU-X1), a Q-switched solid-state 355 nm UV-A laser (Powerchip, Teem Photonics), a C-APOCHROMAT 63x/1.2 W Korr UV-VIS-IR water immersion objective (Behrndt et al., 2012), and a home-made cooling stage were used. The membrane of tail epidermal cells of tailbud embryos was labeled with FM-64 (ThermoFisher). Each ventral midline epidermal cell was cut along the apico-basal axis (5 to 10 μm lines each) by applying 25 UV pulses at 0.7 kHz. The embryos were imaged every 0.2 s frame rate with an exposure time of 150 ms. Single fluorescent images were used to measure tail relaxation 3 s post-ablation, and the percentage of relaxation was calculated as the area of movement of the tail region 3 s after laser cutting.

### Gene knock-down and overexpression

The MOs (Gene Tools, LLC) against *Msxb* and *Admp,* which block translation, were designed according to the previous study (Imai et al., 2006; Waki et al., 2015); *Admp,* 5’-TATCGTGTAGT TTGCTTTCTATATA-3’; *Msxb,* 5’-ATTCGTTTACTGTCATTTTTAATTT-3’. These MOs at 0.25 to 0.50 mM were injected into an unfertilized egg and incubated until observation. To know the phenotype of *Admp* MO embryo at the single-cell level, embryos were stained by Alexa 546 phalloidin (Thermo Fisher).

The DNA constructs used for overexpression of *bmp2/4* under the *Dlx.b* upstream sequence (Ciinte.REG.KH.C7.630497–632996|*Dlx.b*) were used previously (Imai et al., 2012). These DNA constructs were introduced by electroporation.

## Supporting information

Suppl. Mov. 1

Suppl. Mov. 2

Suppl. Mov. 3

Suppl. Mov. 4

**Suppl. Mov. 1. Time-lapse movie of WT (left) and *Admp* MO (right) embryo from late neurula stage 16 to late tailbud stage 25.**

Both embryos are incubated in the same dish. The WT embryo was stained with NileBlue B (Matsumura et al., 2020). The movie frame was surrounded by different colors depending on the tail morphology. red: ventroflexion, blue: relaxation of ventroflexion, yellow: dorsiflexion.

**Suppl. Mov. 2. The z-stack section of the ventral epidermis**

The z-stack images of the ventral midline of the tailbud embryo by phalloidin staining. All the section of the ventral epidermis of the WT and the *Admp* MO embryo was observed at st.22. The F-actin was stained by phalloidin. The arrowheads indicate the TSBC.

**Suppl. Mov. 3. Laser cutting experiment of the midline tail epidermis.**

The dorsal (left) and ventral (right) midline epidermal AP cell borders of the WT mid-tailed embryos were cut with a laser cutter. The arrowheads indicate the cutting point by the laser cutter. The angle of the ventroflexion was relaxed when cut on the ventral side but not on the dorsal side, indicating the AP stress of the ventral midline epidermis (N = 3 each).

**Suppl. Mov. 4. The laser cut of the ventral epidermis with ML and AP direction.**

The ventral epidermis of the mid-tailbud embryo was cut in AP and ML directions with a laser cutter.

**Suppl. Fig. 1. Morphants of tail bending.**

(A) Mid-tailbud stage embryo of WT, DMSO treatment, and dorsomorphin. DMSO and dorsomorphin were treated after the mid neurula stage (st. 15). The dorsomorphin-treated embryo did not bend its tail, similar to the *Admp* MO embryo (Fig. 1A).

(B) Midline section view of WT dorsomorphine-treated and *Admp* MO embryo at st. 18, st. 20, and st. 22 by F-actin staining. The dorsomorphin-treated embryo did not bend its tail (N = 10/10, 12/12 and 9/9), similar to the *Admp* MO embryo (N = 5/5, 13/13 and 15/15).

(C) The phenotype of *Msx-b* MO embryo at st. 22 in one experiment. The ventral tail bending was normally observed. Note that similar phenotypes were reported (Waki et al., 2015).

Abbreviations: A, anterior; D, dorsal; V, ventral; P, posterior

**Suppl. Fig. 2. The asymmetrical actomyosin localization of the notochord.**

(A) F-actin localization of notochord in WT and *Admp* MO was measured on the yellow lines across the dorso-ventral (DV) axis. Abbreviations: A, anterior; D, dorsal; V, ventral; P, posterior.

(B) The intensity ratio between ventral and dorsal F-actin. The ratio is calculated from the peak intensity values in (A) data. If the intensity of the ventral side (red arrow in A) is stronger than the dorsal side (orange arrow in A), the ratio becomes more than 1. In Admp MO embryos, asymmetrical localization remained. F-actin is significantly localized ventrally (WT, N = 8; Admp MO, N = 10). Asterisks indicate statistical significance (t-test, *: p < 0.05). The error bar indicates SD.

**Suppl. Fig. 3. The alignment of the tail midline epidermal cells during the tailbud period.**

From 3D reconstructed confocal stack images by F-actin staining of tailbud embryos (st. 18 in A, st. 19 in B, st. 20 in C, st. 21 in D, st. 22 in E, st. 23 in F and st. 24 in G), cell shapes of both dorsal and ventral midline epidermal cells are traced (yellow lines). Note that cell-cell intercalation completes earlier on the dorsal midline than on the ventral midline.

**Suppl. Fig. 4. pMLC antibody staining during tailbud period.**

(Aa–Hc’) Antibody staining of pMLC from st. 18 to st. 24 in wild type (rows A to G) and st. 22 in Y27632-treated (row H) embryos. Arrowheads show pMLC accumulation in the ventral midline epidermal cells. The part surrounded by the dotted square is enlarged to the panel on the right. Brackets indicate the range of the midline epidermal region. The midline epidermal cells are intercalated, narrowing the region (see Suppl. Fig. 2). Scale bar = 20 μm.

**Suppl. Fig. 5. The alignment of the tail epidermal cells of DMSO-treated and Dorsomorphin-treated embryo at st. 24.**

The cell-cell intercalation was completed in the DMSO-treated embryo (left), and the tail epidermal cells consist of eight rows: dorsal (yellow), two dorsal medio-lateral (green), ventral (red), two ventral medio-lateral (orange), and two laterals (blue) rows. On the other hand, there is a ventral-side specific inhibition of the intercalation (red and orange gradation-colored cells) in the dorsomorphin-treated embryo (right). (N = 2 in DMSO and 2 in dorsomorphin), scale bar: 10 μm.

**Suppl. Fig. 6. Boat cells make up the bending of the tissue.**

(A) Development view of a 3D model schematically showing the boat cell in the ventral midline epidermis (see Fig. 3A–C).

(B) A combination of boat cells, consist of 7 (A), representing the ventral midline epidermis during intercalation. (C) Lateral view of (B). The tissue is bending, as shown in the blue arrow.

**Suppl. Fig. 7. The cell shape change by ML accumulated pMLC during early intercalation.**

In the schematic figure of the apical surface of ventral epidermal cells, the red color indicates the ML localization of actomyosin at the protrusions formed, and the arrows indicate the contractility of actomyosin. This contractility elongates the ventral epidermal cells in the ML direction.

