## Supplementary figures and images for "*Admp* regulates tail bending by controlling ventral epidermal cell polarity via phosphorylated myosin localization"

### Suppl. Fig. 1

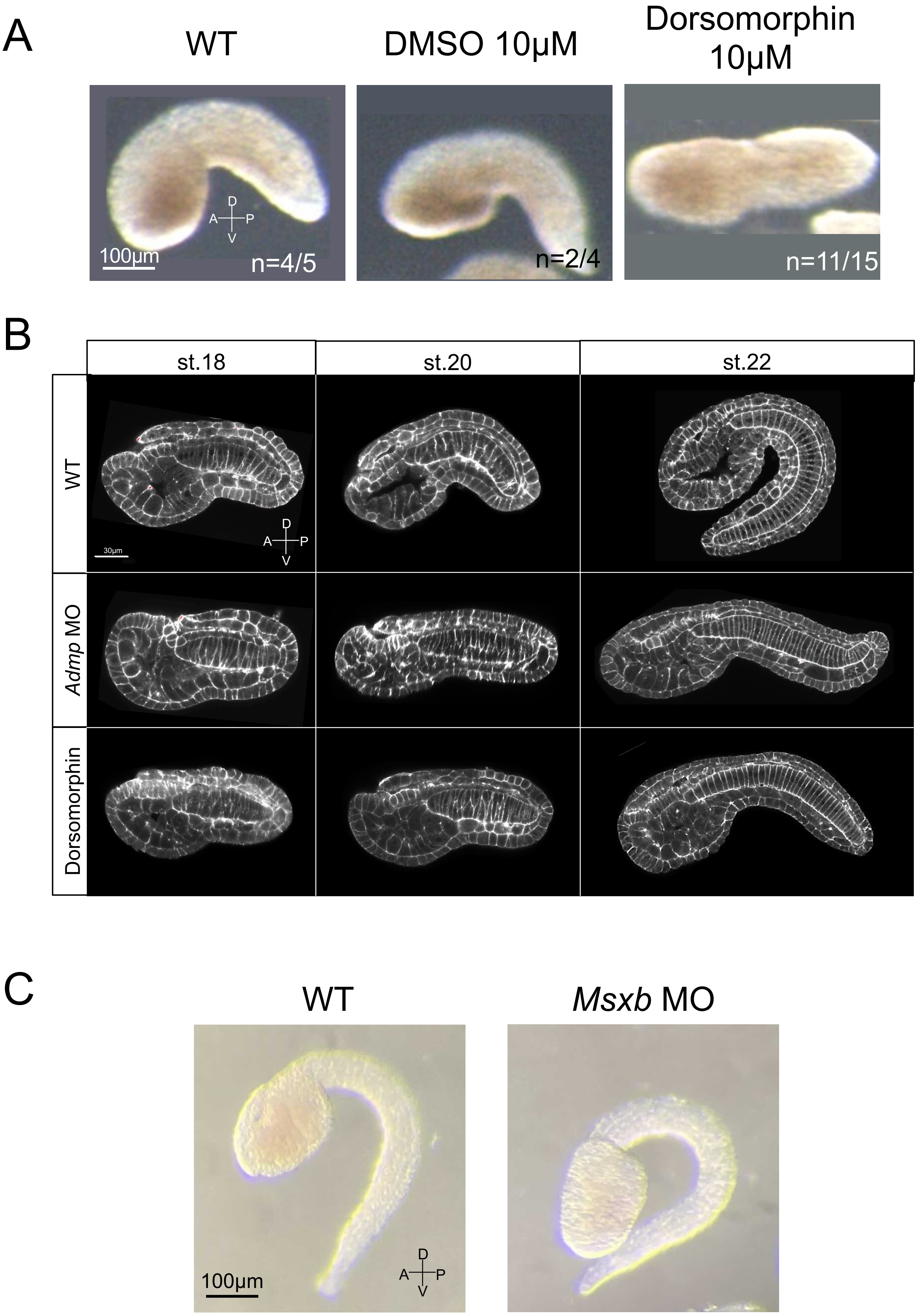

### Suppl. Fig. 2

A

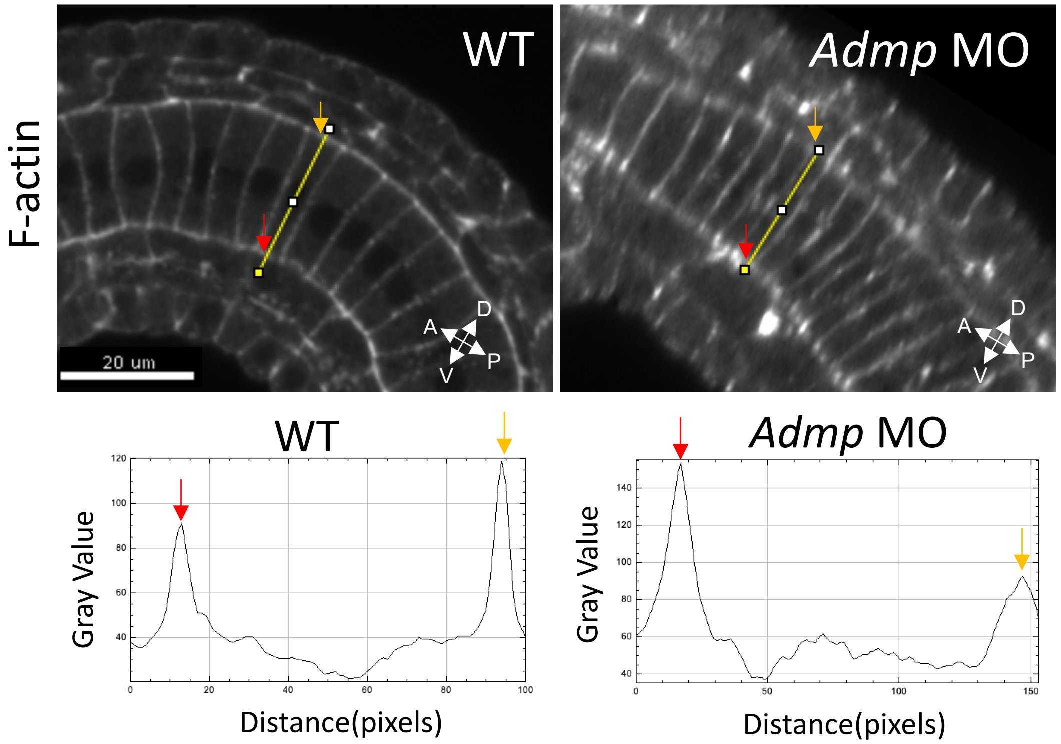


B

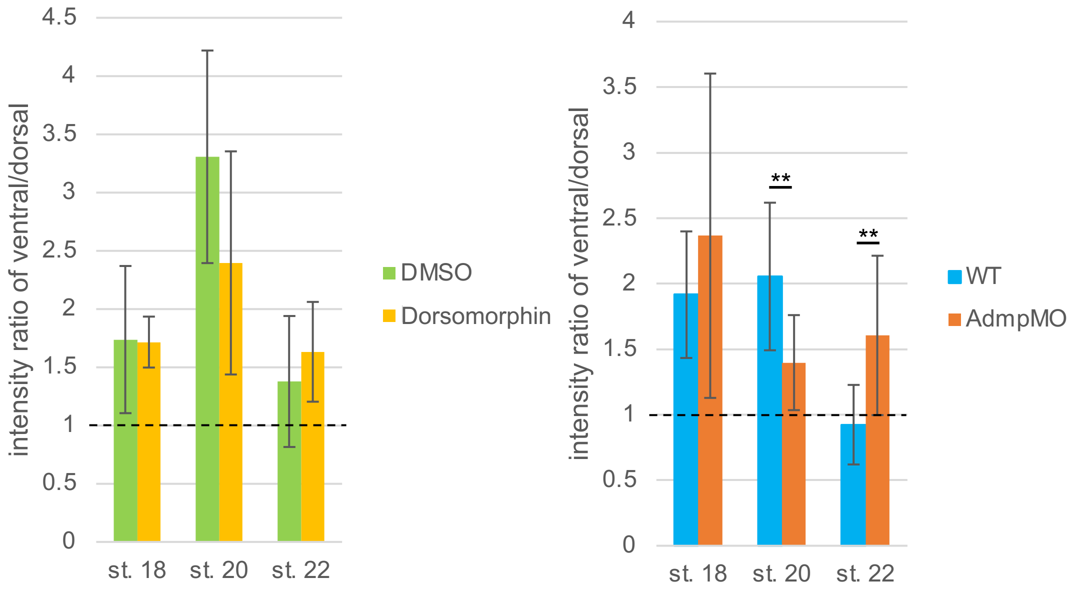


**Suppl.Fig. 2**

### Suppl. Fig. 3

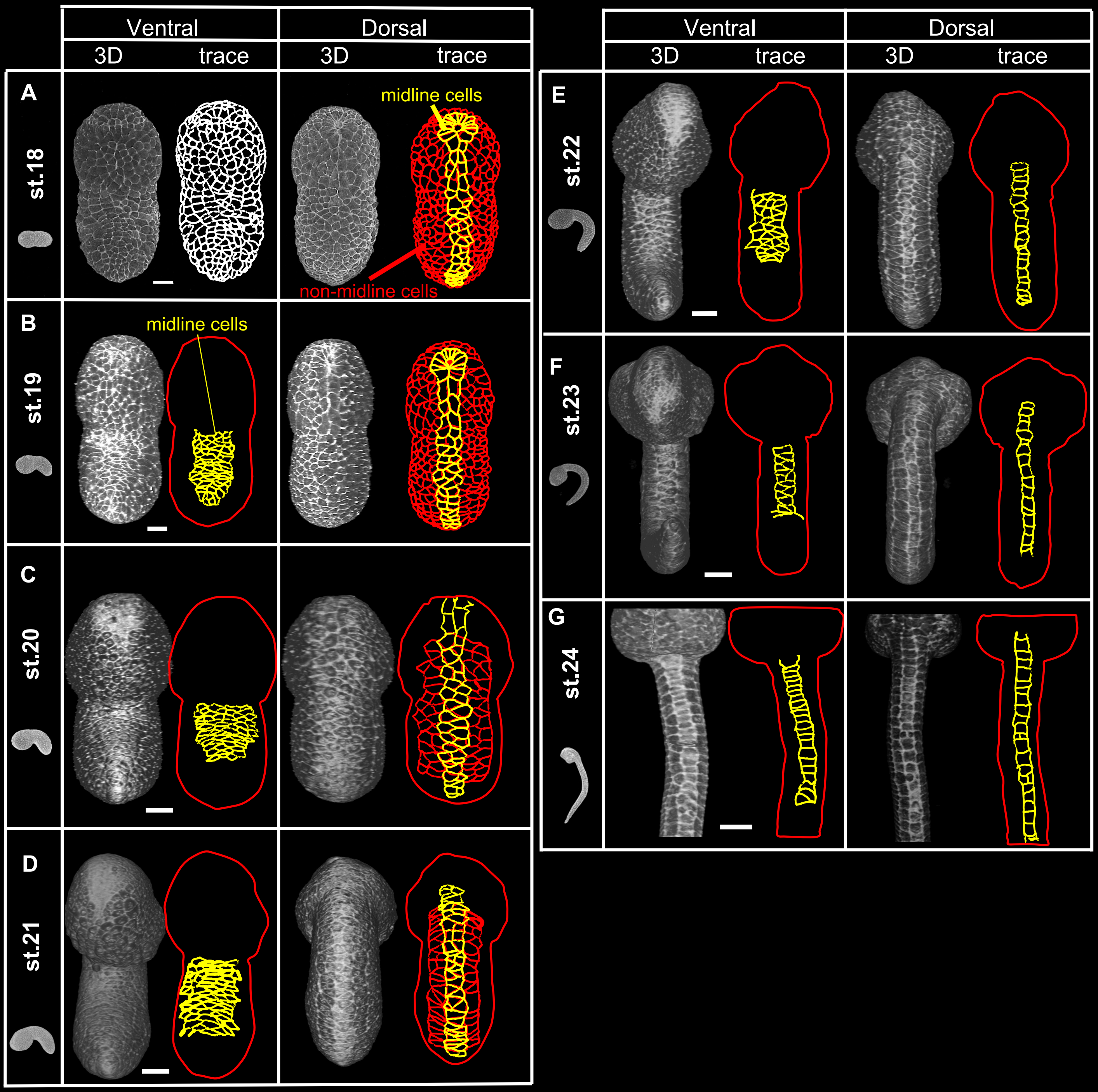

### Suppl. Fig. 4

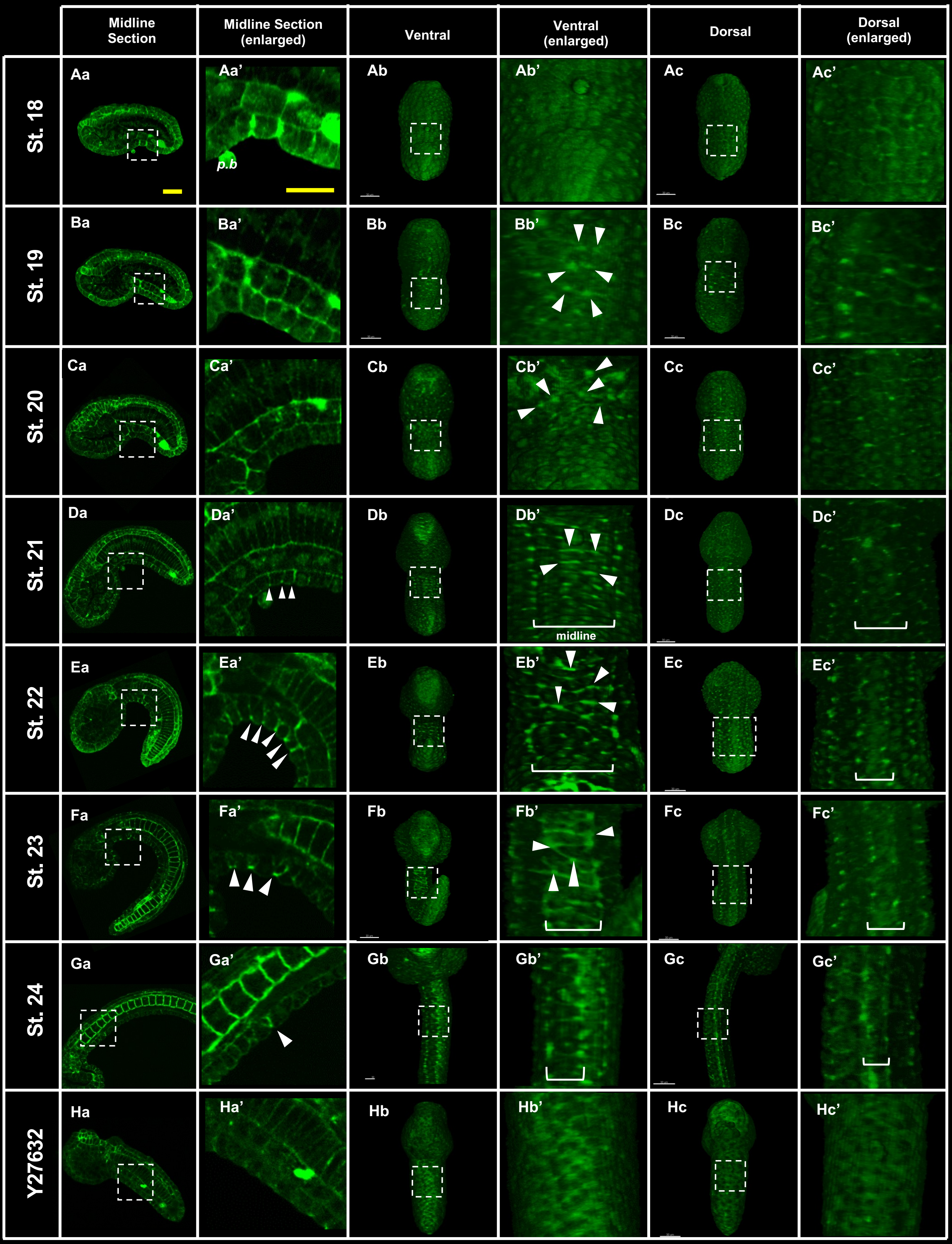

### Suppl. Fig. 5

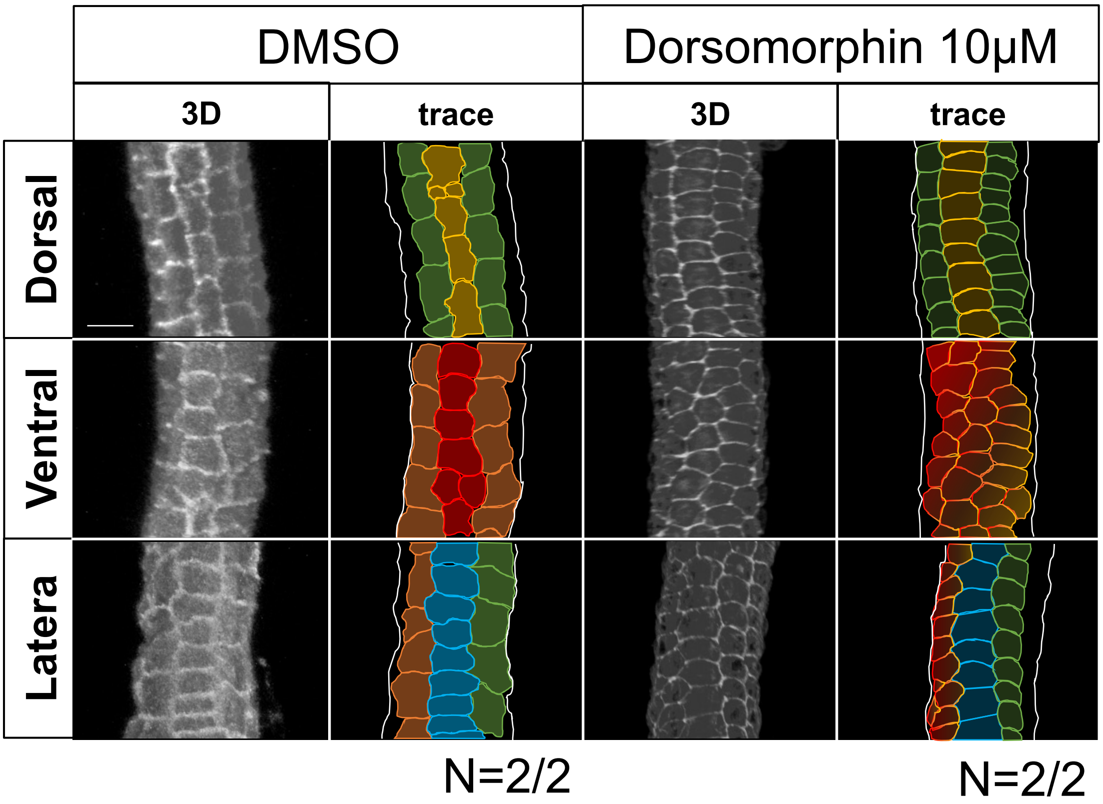

### Suppl. Fig. 6

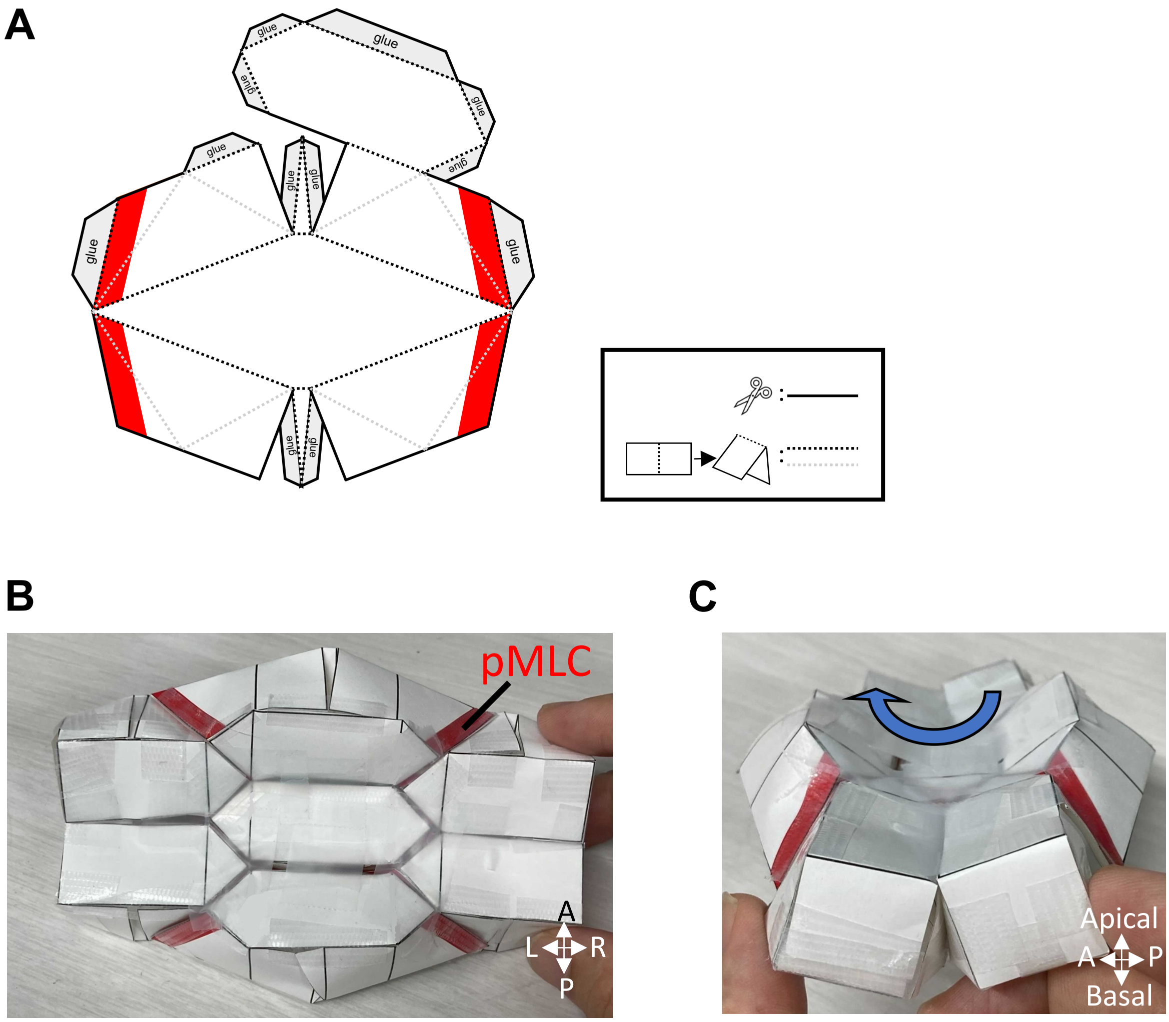

### Suppl. Fig. 7

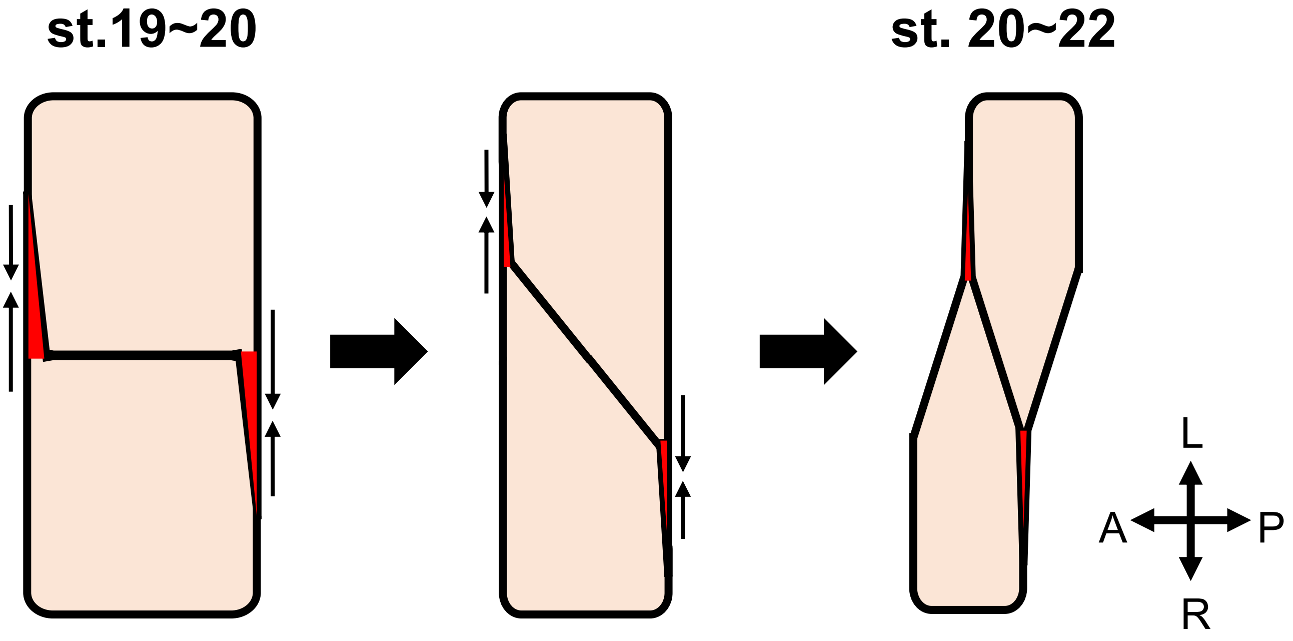


Suppl.Fig. 7
